# Unfolding the hippocampus: an intrinsic coordinate system for subfield segmentations and quantitative mapping

**DOI:** 10.1101/146878

**Authors:** Jordan DeKraker, Kayla M. Ferko, Jonathan C. Lau, Stefan Köhler, Ali R. Khan

## Abstract

The hippocampus, like the neocortex, has a morphological structure that is complex and variable in its folding pattern, especially in the hippocampal head. The current study presents a computational method to unfold hippocampal grey matter, with a particular focus on the hippocampal head where complexity is highest due to medial curving of the structure and the variable presence of digitations. This unfolding was performed on segmentations from high-resolution, T2-weighted 7T MRI data from 12 healthy participants and one surgical patient with epilepsy whose resected hippocampal tissue was used for histological validation. We traced a critical hippocampal component, the hippocampal sulcus and stratum radiatum, lacunosum moleculaire, (SRLM) in these images, then employed user-guided semi-automated techniques to detect and subsequently unfold the surrounding hippocampal grey matter. This unfolding was performed by solving Laplace’s equation in three dimensions of interest (long-axis, proximal-distal, and laminar). The resulting ‘unfolded coordinate space’ provides an intuitive way of mapping the hippocampal subfields in 2D space (long-axis and proximal-distal), such that similar borders can be applied in the head, body, and tail of the hippocampus independently of variability in folding. This unfolded coordinate space was employed to map intracortical myelin and thickness in relation to subfield borders, which revealed intracortical myelin differences that closely follow the subfield borders used here. Examination of a histological sample from a patient with epilepsy reveals that our unfolded coordinate system shows biological validity, and that subfield segmentations applied in this space are able to capture features not seen in manual tracing protocols.

**Research highlights:**

- SRLM in hippocampal head consistently detected with 7T, T2 isotropic MRI
- Hippocampal grey matter unfolded using Laplace’s equation in 3D
- Intracortical myelin and thickness mapped in unfolded coordinate space
- Unfolded subfields capture critical structural regularities and agree with histology

## 1. Introduction

Researchers often distinguish the hippocampus from neocortex but the hippocampus, in fact, also has a cortical composition sometimes referred to as archicortex due to its wide evolutionary preservation (e.g. Duvernoy *et al.*, 2013). Like the neocortex, the hippocampus shows variable gyrification, often referred to as digitations or pseudo-digitations in the anterior hippocampal head and more posterior body/tail, respectively. This creates major challenges for cross-participant alignment and segmentation. This is particularly of interest given the recent controversy over segmentation of the hippocampus into subfields in MR data, which are not sensitive to most cytoarchitectonic features that define the subfields (for an overview of this controversy see Yushkevich *et al.*, 2015a and harmonization efforts by Wisse *et al.*, 2017).

Though present in the rest of the hippocampus, digitations are most prominent in the hippocampal head. This structural feature has created a major challenge for subfield segmentation protocols and as such most protocols do not segment this region, or do not honour its complex and variable structure (see Yushkevich *et al.*, 2015a). Ding & Van Hoesen (2015) recently provided detailed descriptions of the hippocampal head including three different morphologies (2, 3, or 4 digitations), however in our dataset we observed cases with even more digitations that continue through the hippocampal body (see also Gao et al., 2016) and cases with differences in the amount of medial curvature of the uncus. Dalton et al. (2017) and also Berron et al. (2017) have recently published protocols leveraging Ding & Van Hoesen’s descriptions, however, their protocols collapse across different morphologies and deal primarily with one canonical case. This may produce results that are close to the ground truth under different morphologies as well, however, differences in folding will cause a topological shift and so each subfield border should shift in turn. Thus attempting to impose borders without considering topology creates challenges in subjects with different degrees of folding, or different rotations or positions within the medial-temporal lobe (e.g. varying degrees of dysplasia), similar to the challenge of aligning the neocortex in participants with variable gyrification.

In the neocortex, the challenge of inter-subject alignment in cases of variable gyrification have been largely overcome using topology-preserving surface-based alignment (Dale et al., 1999, Fischl et al., 1999a, Fischl et al 1999b, Fischl et al., 2001), which has lead to the development of powerful methods for parcellation (e.g. Glasser et al., 2016). These types of methods have not yet been applied to the archicortex of the hippocampus. However, several studies reported by Bookheimer and colleagues have implemented a technique that is similar but used primarily for visualizing results rather than as an analysis technique (see Ekstrom *et al.*, 2009; Zeineh *et al.*, 2003; and also in 7T MRI Suthana *et al.*, 2015). Under their protocol, delineation of MTL neocortex and hippocampus are performed in the subject’s native space and conformal mapping is used to flatten this tissue such that results can be viewed in a single plane. However, this protocol does not make use of some of the advantageous features used in neocortical surface-based analysis: it does not use a standardized set of coordinate points, segmentation is performed based on a geometric landmarks in native space (i.e. prior to any surface-based alignment), and although the folded topology in the hippocampal body is captured, the digitations and medial curvature of the hippocampal head and tail do not appear to be separately delineated – instead they are labelled using a similar coronal scheme as the hippocampal body. As a consequence, topology is not fully preserved in these areas.

### 1.1 Goals of the current study

The current study aimed to examine the topological structure and ontogeny of the hippocampus in order to develop a two-dimensional coordinate system for alignment and segmentation of variably folded hippocampi across individuals, similar to surface-based alignment methods used in the neocortex. Specific structural features we identified and aimed to account for are the medial folding forming the classic hippocampal C-shape (or inverse C-shape depending on hemisphere and orientation), long-axis and uncal curvature, digitations, and inter-individual variability in each of these features (further detailed in the next section). After tracing each of these features in 7T T2-weighted MR images, we applied the Laplace equation to divide hippocampal archicortex into a set of standardized long-axis and proximal-distal coordinates using anatomically motivated boundary structures that are topologically continuous with the hippocampus. We examined a segmentation of the hippocampus based on the histological samples used by Ding & Van Hoesen (2015) under the framework of this two dimensional, topology-preserving coordinate space, which we then validated by comparison to quantitative MR measures of intracortical myelin and thickness, as well by direct validation in a surgically resected tissue sample from a patient with epilepsy (i.e. comparison of preoperative segmentation to postoperative histology).

### 1.2 Critical structural features we aim to accommodate

During development, the hippocampus originates from a single flat tissue, which in addition to its long-axis curvature, also folds medially upon itself forming a C-shape while differentiating into the various subfields (Duvernoy *et al.*, 2013; Williams, 1995; Smith, 1897)(Figure 1A). This developmental characteristic has several interesting consequences for the structure of the adult hippocampus: all subfields make up adjacent segments of a contiguous tissue segment (though the dentate gyrus makes up a distinct tissue but keeps a consistent position at the distal edge of the CA fields). The sulcus, or ‘crease’ around which this folding occurs can be visualized in histology as the hippocampal sulcus, or in MRI as the ‘dark band’ or SRLM (SLM in the subiculum) after the high myelin laminae thought to be driving its contrast (Kerchner *et al.*, 2010; Thomas *et al.*, 2008, and others), although no extant work has ruled out contributions of non-penetrating blood vessels within the hippocampal sulcus in this contrast. Many subfield segmentation protocols rely on this image feature in the hippocampal body. In the current study we aimed to capture the SRLM in the hippocampal head and tail as well, which we then critically leveraged to differentiate the folds of the entire hippocampus, preserving its topology.

**Figure 1.**
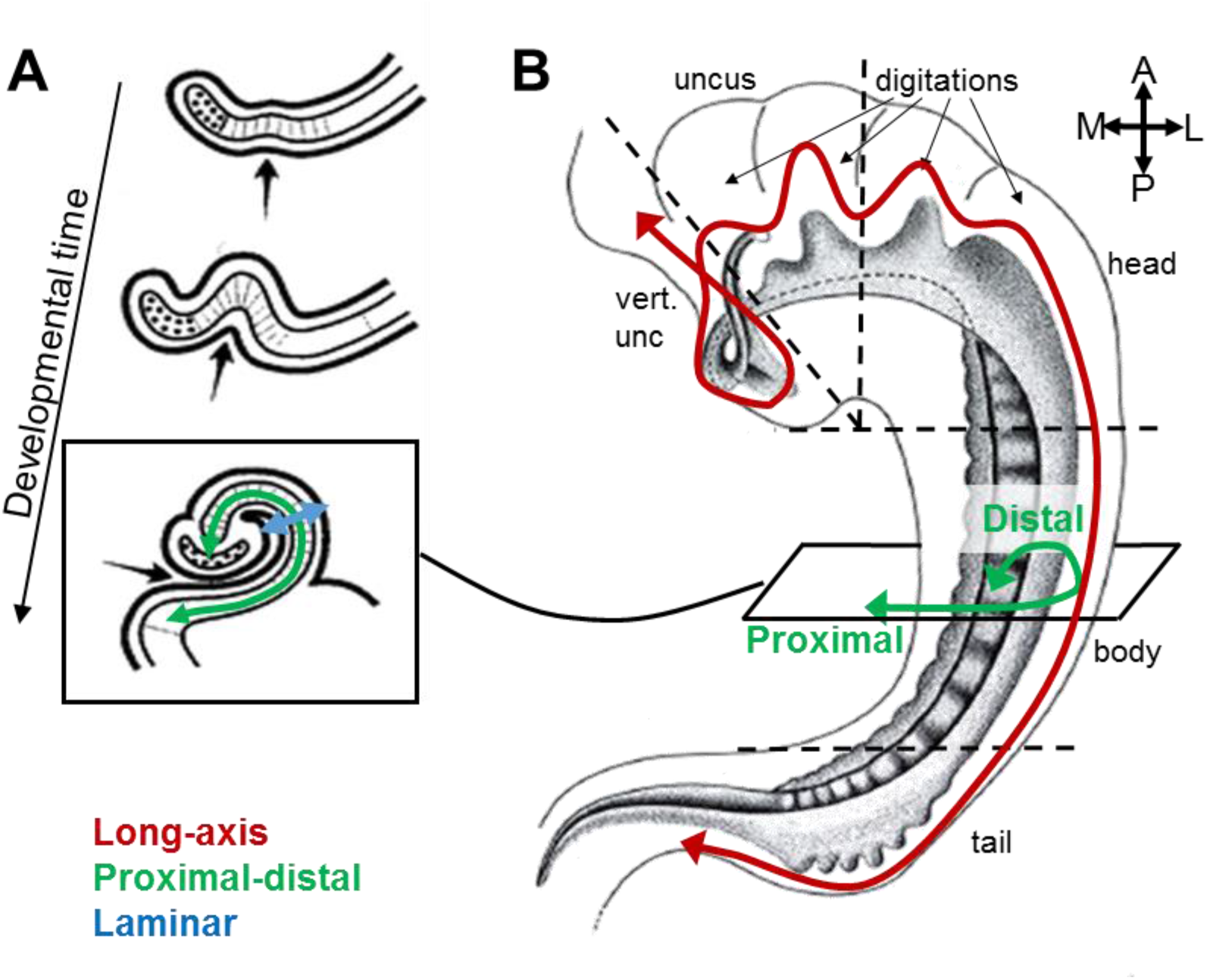
<1.5 column width, coloured>. Ontogeny and anatomy of the hippocampus. A) Coronal sections of the initially flat tissue of the hippocampus folding medially during development, forming the hippocampal sulcus (indicated by black arrow). B) Fully developed hippocampus seen from above, showing the long-axis curvature in the head and tail and anterior digitations. Dotted lines roughly indicate commonly used long-axis divisions of the hippocampus. Both images adapted with permission from (Duvernoy *et al.*, 2013) <permission pending>.

Recent histological evidence from Ding & Van Hoesen (2015) offers a new morphological characterization of the hippocampal head, which we also aimed to respect in our unfolded coordinate space. A main finding in this characterization was the documentation of considerable interindividual differences in digitations (i.e. folding, similar to the gyrification of neocortex) in the hippocampal head, varying from 2 to 4 digitations, with additional pseudo-digitations sometimes found along the lateral and inferior sides of the hippocampal body and tail. Ding & Van Hoesen also delineated the subfields in detail in the uncus-a part of the hippocampal head that curves medially, and then superiorly (see Figure 1B). In line with Duvernoy *et al.*’s (2013) characterization, Ding & Van Hoesen showed that all subfields of the hippocampus contiguously follow this curvature through the hippocampal head and have their natural anterior termination not in the absolute anterior tip of the hippocampus, but rather in the more medial and posterior vertical component of the uncus (Figure 1B, red line terminating in ‘vert. unc’). As the subfields curve into the uncus, their borders also shift such that the subiculum and CA1 move from the inferior side to the lateral, anterior, and finally superior side. A detailed segmentation of this region must capture each of these features, and here we aim to provide a tool with sufficient validity and precision to index these structural complexities.

## 2 Methods

### 2.1 Study participants

For MRI data acquisition, healthy participants were recruited from Western University, London, Canada (n = 12; 6 females; ages 20-35, mean age 27.6). This study was conducted with Western’s Human Research Ethics Board approval, and informed consent was collected from each participant prior to participation.

### 2.2 MRI acquisition

Imaging was performed using a 7T neuroimaging optimized MRI scanner (Agilent, Santa Clara, CA, USA/ Siemens, Erlangen, Germany) employing a 23-channel transmit-receive head coil array constructed in-house. Four T2-weighted turbo spin echo (TSE) 3D (3D sagittal, matrix: 260x366, 266 slices, 0.6mm^3^ isotropic, ~8.5 mins per scan) images were acquired from each participant. All images were acquired in sagittal rather than coronal oblique orientation for optimal whole brain coverage, given that these data were also used for other, whole brain studies. The use of isotropic acquisition differs from most hippocampal imaging protocols that acquire thick coronal slices oblique to the hippocampus to maximize in-plane resolution. However, these protocols limit the visibility of structures which run perpendicular to the long-axis of the hippocampus, including most of the hippocampal head and tail. By using isotropic voxels, we were able to capture small features such as the hippocampal SRLM in high detail, throughout the entire hippocampus. A T1-weighted MPRAGE (3D sagittal, matrix: 256x512, 230 slices, 0.75mm^3^ isotropic) image was also collected.

### 2.3 Preprocessing

All scans were processed as follows: the first T2-weighted image (scan 1) was upsampled to 0.3mm^3^ isovoxels using cubic spline interpolation, then scans 2, 3, and 4 were rigidly registered it using FSL FLIRT registration (Jenkinson, 2002; Tofts, 2005). All four scans were then averaged together to produce a single, 0.3mm^3^ isovoxel, high-contrast volume. This volume was then reoriented to an oblique orientation, with coronal slices perpendicular to the long axis of the hippocampus, by rigid registration to an average template in coronal oblique orientation (as determined by visual alignment of the long-axis of the template hippocampus to the anterior-posterior axis). T1-weighted scans were registered to this high-contrast coronal oblique T2-weighted volume using rigid registration as above.

### 2.4 Detection and labelling of the SRLM and hippocampal grey matter

Under our isotropic MR acquisition protocol, the SRLM was visible in the entire long-axis of the hippocampus, including the digitations and uncus of the hippocampal head. Exemplary slices from the hippocampal head and body and a 3D model reconstruction can be seen in Figure 2A. All manual tracing was performed in ITK-SNAP 3.4 (Yushkevich *et al.*, 2006), and the built-in ‘Snake’ tool was also used to facilitate tracing. The detailed protocol for tracing, ‘feathering’, dilation, and manual adjustments can be found in Supplementary Materials 1.

**Figure 2.**
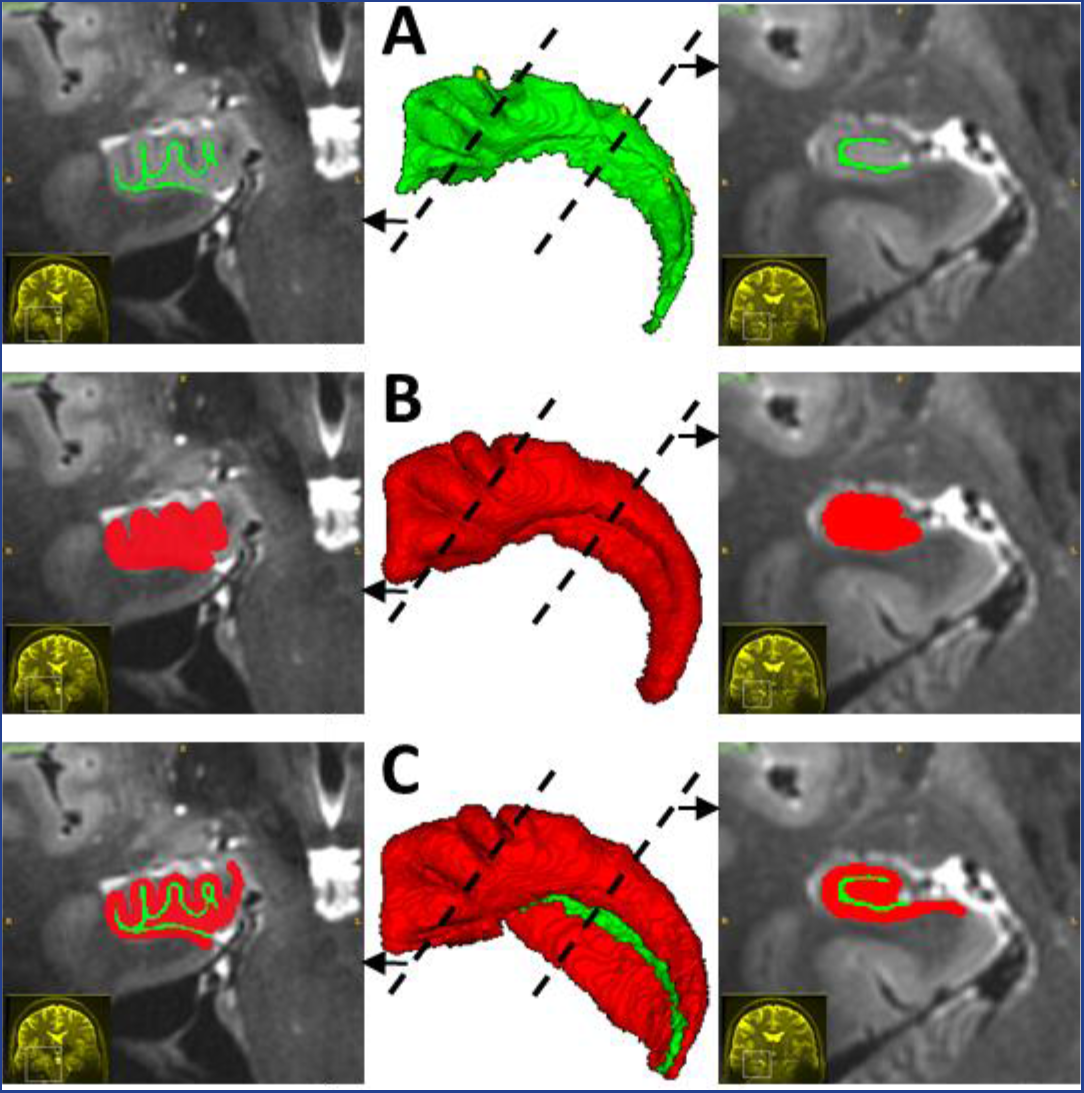
<single column width, coloured>. Illustration of SRLM (green) and hippocampal grey matter (red) labelling. A) 3D model of the SRLM label in the center, with example coronal slices from the head and body on the sides, at the positions indicated by the dotted lines. B) Same views as above, but depicting the SRLM label after spherical dilation. C) Same views as above showing the combined SRLM label and grey matter label, after manual adjustments to the grey matter label.

Much of the morphology of the hippocampus is systematically related to the SRLM. Much of the morphology of the hippocampus is systematically related to the SRLM. This can be observed in our dataset (e.g. Figure 2) where models of the SRLM capture the same digitation and curvature structures as models of hippocampal grey matter, and this also agrees with anatomical descriptions wherein all laminae of the CA fields (including SRLM) and even the underlying dentate gyrus follow the hippocampus’ digitated structure (see Figure 1; Duvernoy et al. 2013). The various subfields of the hippocampus surround the SRLM, and thus we made use of this proximity to initialize grey matter segmentation with the SRLM segmentation. We segmented the gray matter of the hippocampus initially by dilation of the SRLM label using an initial active contour evolution of the SRLM, followed by manual correction. We used ITK-SNAP’s Snake tool (Yushkevich *et al.*, 2006), which evolves a seed region in 3D to fill a structure of interest. In our case, we initialized the evolution using the SRLM and first applied no constraints on the evolution, resulting in uniform, spherical dilation. The amount of dilation was determined by visually inspecting whether the outer borders of hippocampal grey matter had been reached, and varied slightly between traces depending on the available image information. Evolution constrained by edge attraction (with parameters defined by the user based on image quality) and manual adjustments were then applied. We did not trace the SRLM along the superior side of the subiculum (the most medial, less folded extension of the hippocampus) as it was not consistently visible. Accordingly, grey matter was not labelled as part of the dilation of the SRLM label but had to be labelled manually by the rater, using a spherical paintbrush. This was also the case in the most medial, vertical component of the uncus where the SRLM was often not visible. Further manual adjustments included the removal of grey matter label from the CSF on the medial side of the dentate gyrus, and minor adjustments throughout to ensure all grey matter was labelled as such. Because errors in manual segmentation can produce distortions in the next step of hippocampal grey matter unfolding, the unfolding results of each hippocampus were visually examined by the raters to ensure their labelling followed the rules outlined in Supplementary Materials 1.

To assess how reliably the SRLM could be segmented in our high-resolution images, we repeated the segmentation with an additional trained rater, and calculated the spatial overlap between these segmentations using the Dice similarity index (DSI). DSI represents the proportion of overlapping voxels in two segmentation labels over the mean number of voxels per label. It can vary from 0 to 1, with values close to 1 denoting high overlap (Dice, 1945).

### 2.5 Manual subfield segmentation

Before unfolding of hippocampal grey matter, we performed manual segmentation of this tissue into subfields in a set of 10 hippocampi (5 participants x2 hemispheres) with varying numbers of digitations and varying curvature in the uncus. This was done by carefully matching coronal views in our MR images with the closest corresponding histological segmentations in the hippocampal head provided by Ding & Van Hoesen (2015). In the hippocampal body and tail, segmentations were performed based on the descriptions of Duvernoy *et al.* (2013). Example slices of these segmentations can be seen in Figure 3, and additional examples as well as qualitative descriptions can be found in Supplementary Materials 2. Note that it was not our intention to develop a manual segmentation protocol in this paper; we simply aimed to determine whether the trajectories of the hippocampal subfields in our unfolded coordinate space could be captured in a way that respects the recently elucidated complexity in the hippocampal head. Thus, we assessed the reproducibility of these segmentations in only a small sample (four left and four right) using DSI scores as above, to ensure values were at least comparable with previous reports (results in Figure 6).

**Figure 3.**
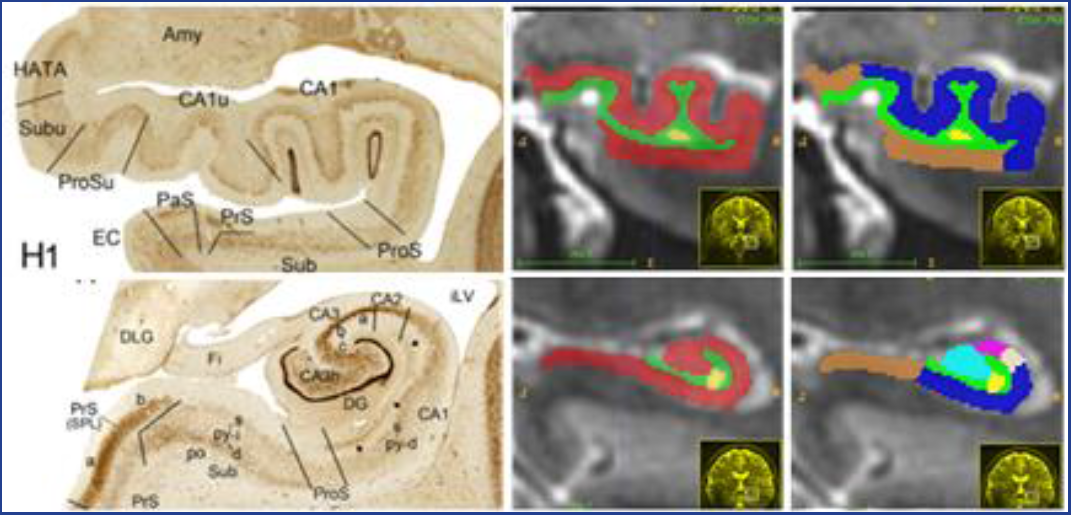
<single column width, coloured>. Example slices showing manual subfield labelling from hippocampal head (top) and body (tail). Left shows reference materials from Ding & Van Hoesen (2015), middle shows manually traces SRLM (green), hippocampal grey matter (red) and cysts (yellow), right shows manually delimited subiculum (brown), CA1(blue), CA2 (white), CA3 (pink), and dentate gyrus (cyan). See Supplementary Materials 2 for further details.

### 2.6 Unfolding of hippocampal grey matter

In the neocortex, 3D computational tools such as Laplace’s equation have been used to precisely and flexibly calculate neocortical thickness (e.g. Jones *et al.*, 2000; Sowell *et al.*, 2004). In principle, the Laplace equation, *∇^2^φ = 0,* defines a potential field (*φ*) whose values change based on their distance from two boundary surfaces. The solution is twice-differentiable *(∇^2^ = 0*), which guarantees a level of smoothness that is appropriate for brain anatomy. In studies of neocortical thickness the potential field spans the neocortical grey matter, while the boundaries are the white matter and pial surfaces. Thickness is then computed by generating streamlines across the resulting potential field gradient.

We reasoned that Laplace’s equation may also be used for unfolding of hippocampal grey matter, not only to determine thickness along the laminar dimension, as above, but also to compute potential field gradients along the long-axis and proximal-distal dimensions. To do so, it is critical to employ multiple sets of boundary conditions, sometimes referred to as ‘source’ and ‘sink’. For example, unfolding along the long-axis dimension makes use of anatomically motivated boundaries at the anterior (source) and posterior (sink) ends of the hippocampus. The potential field in between is defined over all grey matter, and increases smoothly from source to sink. We thus solved Laplace’s for three different equations, *∇^2^φlong-axis = 0*, *∇^2^φproximal-distal = 0*, *∇^2^φlaminar = 0*, to determine a different potential field for each hippocampal dimension. The domain was identical for each dimension, i.e., the hippocampal gray matter, but the boundary conditions were distinct in each of them, defined as anatomical landmarks along at the edges of the hippocampal tissue. An iterative finite-differences approach was used to obtain the solution for each Laplace equation, while employing a 26-neighbour average to compute the updated potential field, and terminating when the potential field change is below a specified threshold (sum of changes < 0.001% of total volume). It is important to note that the SRLM voxels were not included in the gray-matter domain of Laplace’s equation,. This protocol feature effectively provides a barrier such that the streamlines follow a geodesic (i.e. along the outer surface) path along the hippocampus and do not introduce short-circuits. Solving of the Laplace equations was performed in MATLAB (code available at https://github.com/jordandekraker/HippUnfolding).

#### Long-axis dimension boundaries

As discussed in section 1.2, each of the subfields has its natural anterior terminus in the vertical component of the uncus (Ding & Van Hoesen, 2015). Hippocampal grey matter in this area borders the grey matter of the amygdala, making up an area that is typically referred to the hippocampal-amygdalar transition area (HATA) (Ding & Van Hoesen, 2015). At the tail of the hippocampus, a structure named the indusium griseum (which is actually a vestigial extension of the dentate gyrus) extends medially and posteriorly from the hippocampus and then curves upward and anteriorly along the midline of the brain before merging with the cingulate cortex (Duvernoy et al., 2013). The HATA and indusium griseum thus make up two visible structures which correspond to the natural anterior and posterior termini of each of the hippocampal subfields. We manually traced these structures only where they border hippocampal grey matter (see Supplementary Materials 1 for details), and used them as source and sink regions in Laplace’s equation (see Figure 4A for illustration).

**Figure 4.**
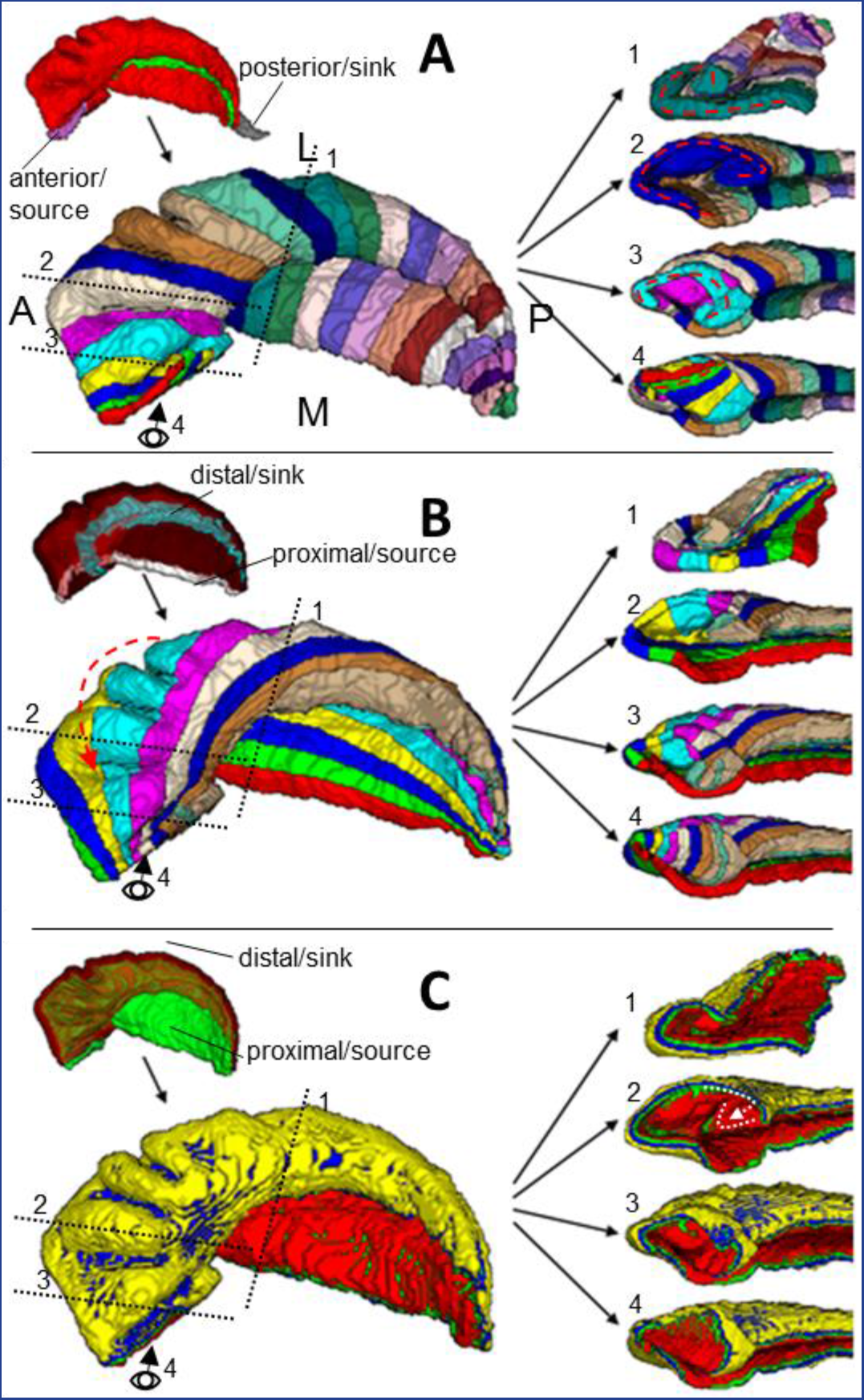
<single column width, coloured>. Illustration of Laplacian unfolding along the long-axis, proximal-distal, and laminar dimensions in A), B), and C), respectively. The upper left inset image in A) shows a 3D model of the SRLM (green) and grey matter (red) labels, with the HATA (pink) and indusium griseum (grey) to be used as boundaries for Laplace’s equation. The lower left image in A) shows arbitrarily coloured bins within the resulting potential field gradient. To the right is the same model as the lower left, but showing cross sections from the body (top) and head (lower three), depicting in particular the uncus (lower two) and vertical component of the uncus (bottom). The locations of these cross sections are shown by the black dotted lines (1-3) and the eye figure (4). B) shows the same views of the same hippocampus, but using the inner dentate gyrus (turquoise) and medial temporal lobe cortex border (white) (upper left insert in B), as boundaries for Laplace’s equation for the proximal-distal dimension. C) shows the same views of the same hippocampus, but using the SRLM (seen in green under semi-transparent red grey matter) and outer hippocampal borders (upper left insert in C) as boundaries for Laplace’s equation for the laminar dimension. White dotted lines in C) (right) show the true laminar structure of CA4 and DG, which is not respected in our laminar potential field gradient.

#### Proximal-distal dimension boundaries

We defined the proximal border as the point at which the subiculum, the most proximal subfield, contacts the grey matter of neighbouring medial-temporal lobe neocortex (see Supplementary Materials 1 for details). To index the full extent of hippocampal grey matter, its distal border can be defined as the granule cell layer of the dentate gyrus (i.e. the part of dentate gyrus which most closely borders the SRLM). We employed a custom approach to detect this tissue: within each previously computed long-axis bin we applied volumetric fast marching (Sethian, 1999) along hippocampal grey matter, starting at the border with surrounding temporal lobe cortex, and estimated the dentate gyrus as being the most distal 12% of this distance (determined experimentally). To index only the innermost granule cell layer of this tissue, we dilated the SRLM by a single voxel (8 nearest neighbours) over this rough dentate gyrus approximation. The result included only the most distal portions of the dentate gyrus, corresponding roughly to the granule cell layer.

An additional challenge in the proximal-distal unfolding of the hippocampus lies in the vertical component of the uncus. Here, hippocampal grey matter, unlike in the rest of the hippocampus, does not follow the classic folded C-shape, and instead flattens out (Figure 4 bottom right). The dentate gyrus persists on the medial edge of this region, but it is not separated by the other subfields with any visible SRLM at the current image resolution. We thus defined the dentate gyrus’ location manually for this region (Figure 4B, third and fourth right panels, turquoise bin).

#### Laminar dimension boundaries

We defined the sink for Laplace’s equation as the outermost surface of the hippocampus, and the source as the SRLM. However, as mentioned in section 2.1, the subiculum and vertical component of the uncus do not border the SRLM. Therefore, they were artificially extended over these regions. For the subiculum, this was performed computationally by dilating the SRLM label along the surface of the subiculum until the most medial point was reached in each coronal slice. For the vertical component of the uncus this label was created manually. These artificially extended labels were used in as the source in Laplace’s equation (Figure 4C).

### 2.7 Subfield borders in unfolded coordinate space

The long-axis and proximal-distal potential field gradients together make up a 2D coordinate system that can be used for indexing columns of hippocampal grey matter. Using this ‘unfolded coordinate space’, the location of each subfield border can easily be indexed. We used the manual subfield segmentations performed on 10 hippocampi (see section 2.5) to generate a subfield atlas in the unfolded coordinate space. That is, for each manually segmented hippocampus, we identified the long-axis and proximal-distal coordinates that correspond to each of the subfield borders. We then averaged these borders together at each long-axis point, and plotted the labelled data in the 2D unfolded coordinate space (see Figure 7).

Given the low variability of the subfield borders in unfolded space, we then applied our Laplacian unfolding to the remaining hippocampi and, rather than performing subfield segmentation manually, we applied the group-averaged borders from Figure 7. That is, for each long-axis and proximal-distal coordinate in each hippocampus, we assigned the corresponding label from Figure 7. We then assessed the overlap of the 10 manually segmented hippocampi to their unfolded group-average border segmented counterparts using Dice Similarity Indices. To avoid bias, we used a leave-one-out approach where a given participant’s manual segmentations (both left and right) were not included in unfolded group-averaged borders.

### 2.8 Quantitative unfolded tissue properties

Properties such as intracortical myelin content and cortical thickness have been shown to be useful for parcellation of the neocortex into functional subregions (e.g. Glasser *et al.*, 2014; Glasser and Van Essen, 2011). The ratio of T1-weighted over T2-weighted values produces a map that is correlated with quantitative R1, and is used as a surrogate measure of intracortical myelin (Glasser and Van Essen, 2011). We estimated intracortical myelin in this way, and estimated cortical thickness by fitting streamlines to the laminar potential field gradients of all hippocampi. We then plotted these values across unfolded coordinate space. To avoid confounds from partial voluming, we mapped the myelin contents of hippocampal tissue from only the middle 25-75%, as determined by our laminar Laplacian field, corresponding to approximately 2 voxels at the current resolution (similar results were obtained when the superficial and deep laminae were included as well). To illustrate how these values map onto the native 3D space of the hippocampus, we generated 3D hippocampal models with surface colouring that corresponds to the underlying myelin estimates from the group average in two hippocampi (one highly digitated and one less digitated exemplar).

### 2.9 Histological validation

One temporal lobe epilepsy patient with left mesial temporal sclerosis (age 34; male) underwent preoperative 7T scanning and then went on to receive a left anterior temporal lobectomy, with inclusion of the amygdala and hippocampus, as part of their standard of care. The surgical tissue underwent a standardized protocol involving overnight scanning in a ultra-high field *ex vivo* 9.4T MRI, agar embedding, and cutting into blocks 4.4mm apart for paraffin embedding and histological sectioning. Staining with H&E, Neu-N, GFAP, and Luxol fast blue was performed, and slides were digitized at 0.5 micron/pixel resolution. The subfields were manually annotated on the Neu-N histology images by rater KF using the Aperio ImageScope software, with criteria outlined in Ding & Van Hoesen (2015) and verified by experienced pathologist, Dr. Robert Hammond (see Supplementary Materials 3 for additional details on how these segmentations were performed). We employed our previously developed and validated pipeline for MRI and histology registration (Goubran *et al.*, 2015, 2013) to perform direct validation of 7T hippocampal subfield segmentation against ground-truth histological sections. The histology-MRI image registration procedure involved iterative 2D-3D deformable registration of downsampled (100 micron/voxel) histology slides to the reference 9.4T tissue MRI, along with 3D deformable landmark-based registration of the 7T MRI to the 9.4T MRI. Segmentation labels as well as the proximal-distal gradient from the *in vivo* 7T images were then propagated to this aligned histology space for direct comparison.

## 3 Results

### 3.1 Detection and labelling of the SRLM and hippocampal grey matter

To assess reliability, inter-rater DSI was calculated for SRLM and grey matter labels. Note that DSI tends to be lower for thin structures at higher resolutions because of the high surface area to volume ratio. DSI revealed good spatial overlap in both the SRLM (0.72±0.03 right; 0.70±0.04 left) and hippocampal grey matter (0.84±0.01 right 0.81±0.02 left). Thus, our dataset contained sufficient contrast to detect and label the SRLM and grey matter based on the visual features described in Supplementary Materials 1 with good consistency.

### 3.2 Subfield borders in unfolded coordinate space

Hippocampal subfields projected into unfolded coordinate space are shown in Figure 5. As expected, the same proximal-distal arrangement of subfields was found throughout the entire hippocampus in unfolded coordinate space, including the hippocampal head (Figure 5). Variability was low for all borders (i.e. low SEM compared to the area of each subfield), and no subfields crossed over each other either in the group average or in any given unfolded segmentation example. Reliability of the manual subfield segmentations (based on Ding & Van Hoesen, 2015) and comparison of unfolded group-average segmented hippocampi to their manually labelled counterparts are shown in Figure 6. Lowest DSI was seen for the smallest subfields-CA2 and CA3. The data show that combining these two labels, as in some other manual segmentation protocols, leads to moderate improvements.

**Figure 5.**
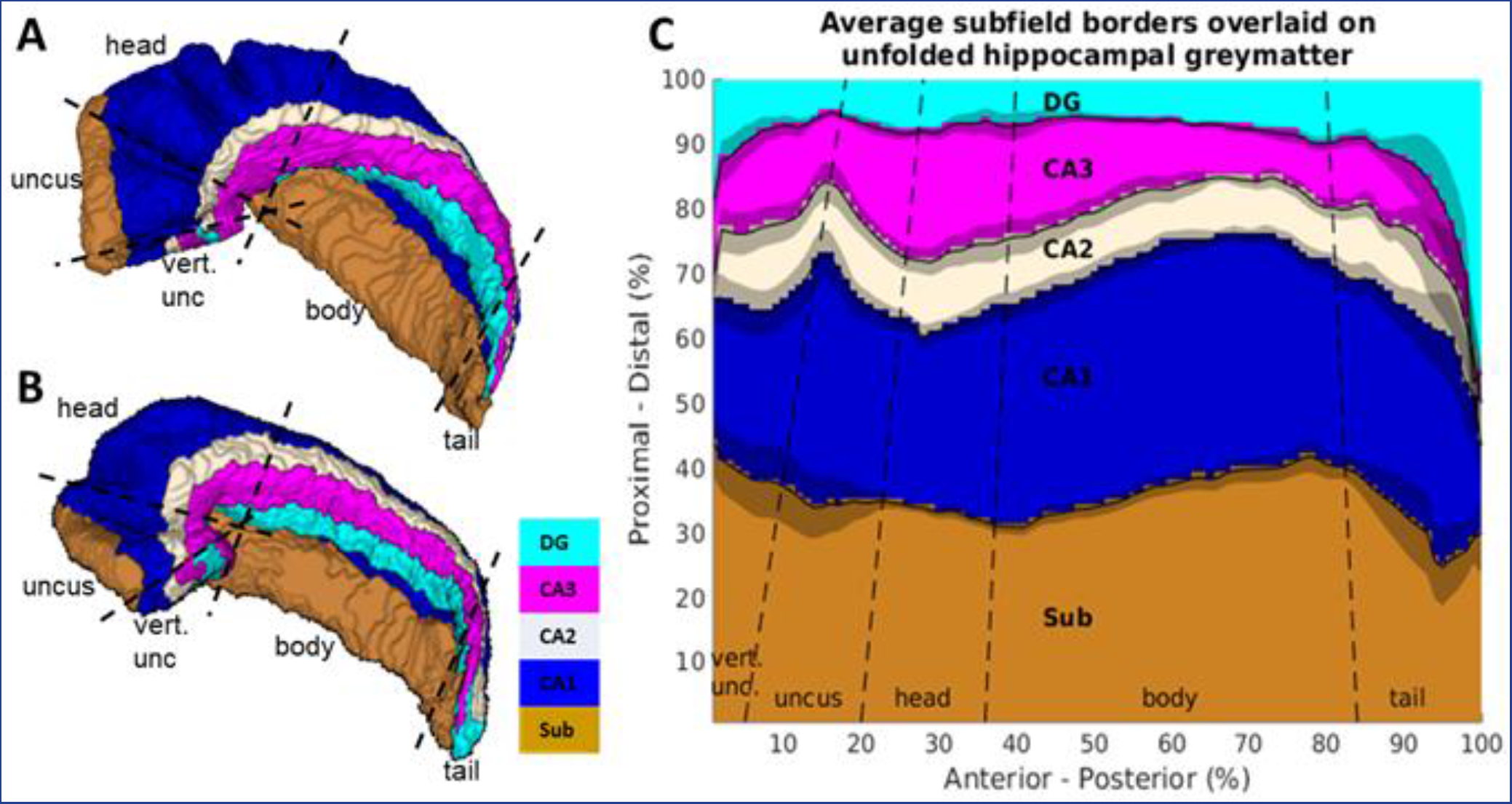
<double column width, coloured>. Hippocampal segmentations in unfolded coordinate space. A) Example of a manual subfield segmentations based on (Ding & Van Hoesen, 2015). B) Same as A), but showing an exemplar with less digitations and medial curvature. C) Flattened out or ‘unfolded’ hippocampal grey matter, with subfield label identity determined at each long-axis and proximal-distal coordinate from the manual segmentations (winner-takes-all over the sample). The shaded areas indicate standard error of the mean for each subfield boundary location across the sample of manual segmentations. Dotted lines approximately indicate commonly used boundaries between the hippocampal head, body, and tail, with the head further subdivided into uncus and vertical component of the uncus.

**Figure 6.**
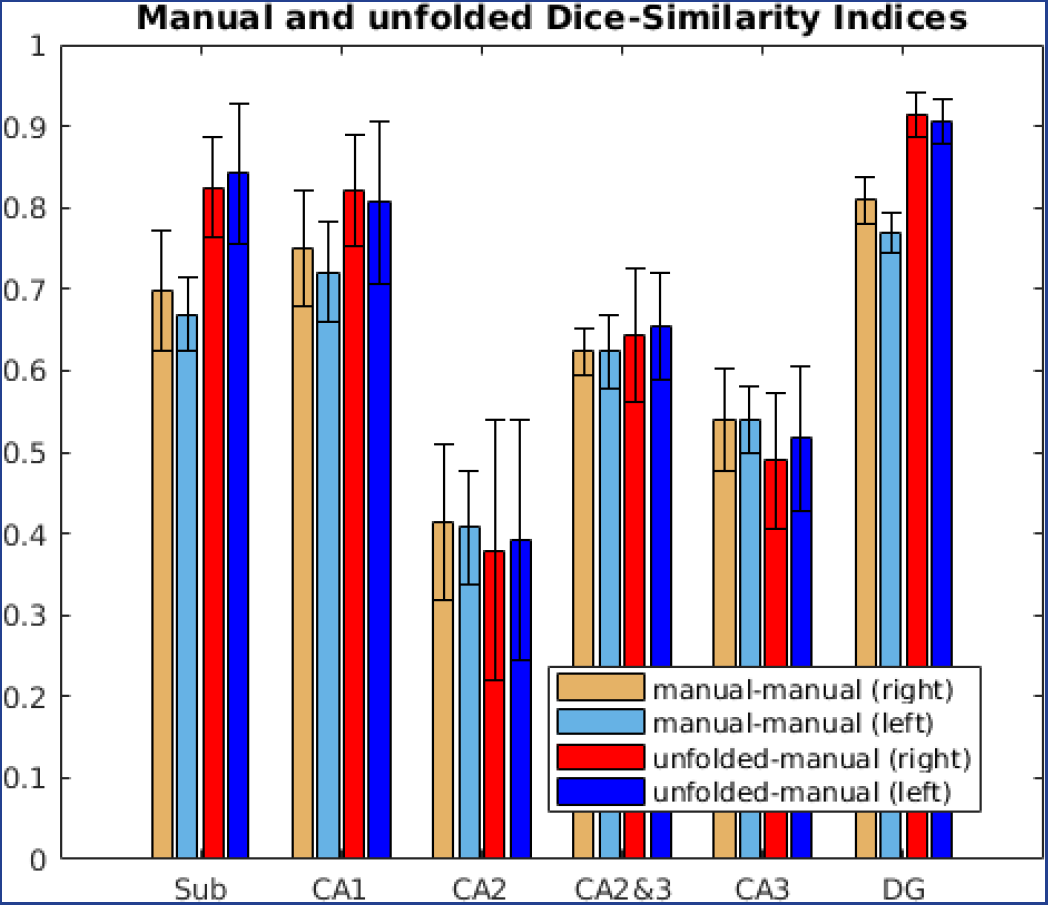
<single column width, coloured>. Spatial overlap in Dice Similarity Index (DSI) between manual subfield segmentations (manual-manual) and between leave-one-out unfolded group-average subfield segmentations and their manually segmented counterparts (unfolded-manual). The leave-one-out technique was performed such that borders from one participant’s left and right hippocampi were not included in the averaged borders that informed unfolded segmentation of that participant’s hippocampi.

### 3.3 Quantitative unfolded tissue properties

Unfolding provides a way to view grey matter properties across the entire extent of the hippocampus in a single 2D view. This unfolded view can obviate patterns that are not apparent when limited to single slices in native 3D space. Here, we mapped intracortical myelin and cortical thickness (Figure 7A and B, respectively). It should be noted that additional properties, including those used by Glasser *et al.*, (2016), can be mapped in this way as well. We additionally mapped these results to the surface of an example 3D hippocampal model in order to visualize them in native space and to allow for easier comparison with manual and unfolded average segmentations (Figure 7C).

**Figure 7.**
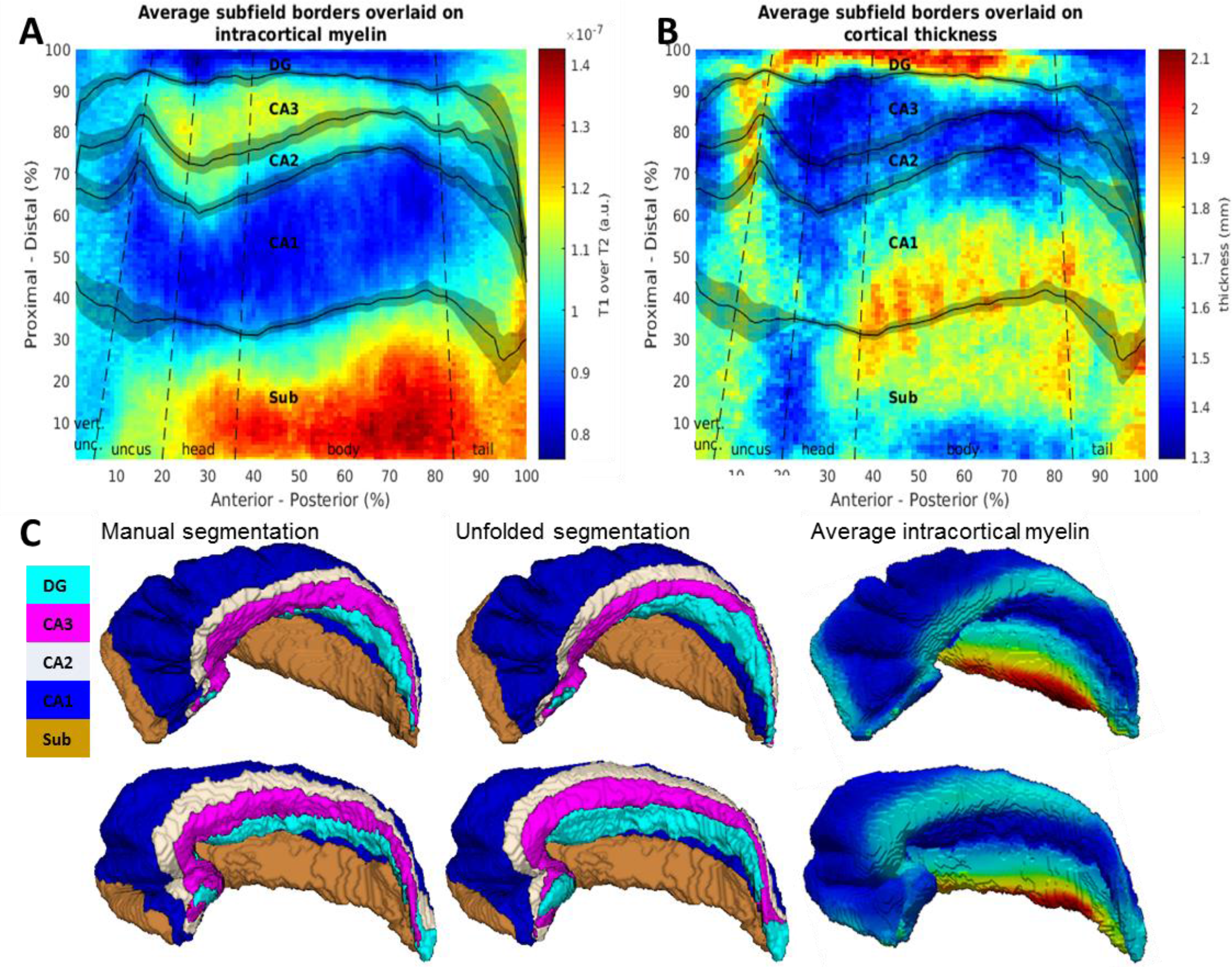
<double column width, coloured>. Quantitative mapping in unfolded coordinate space compared to subfield borders. A) Average intracortical myelin estimates (T1 over T2 MR intensities; arbitrary units). B) Average cortical thickness. Both A) and B) have average subfield borders overlaid. Note that in the dentate gyrus, thickness estimates are actually perpendicular to the true laminar structure (see section 3.2). C) Manual and unfolded subfield segmentations compared to intracortical myelin in highly digitated (top) and less digitated (bottom) example hippocampal models. Average intracortical myelin is mapped to the surface models of these hippocampi for easier comparison.

### 3.4 Histological validation

Segmentation on *in vivo* 7T data from the surgical patient are compared to the same patient’s *ex-vivo* resected and histologically stained hippocampus in Figure 8. Atrophy and cell loss in area CA1, CA3, and the dentate gyrus, with relative sparing of CA2, can be seen in the epileptogenic tissue (Figure 8, far left). Cell loss is most evident in the distal portions of CA1 where reduced staining is seen, and atrophy is apparent from the small area occupied by each subfield compared to histological references (e.g. Ding & Van Hoesen 2015) and reduced digitation structure (see Oppenheim *et al.*, 1998). These findings describe classical hippocampal sclerosis (Blümcke *et al.*, 2007). Note also that this patient shows only two clear digitations in the hippocampal head, which were not well captured in the *in-vivo* labelling of grey matter tissue (i.e. digitations cannot be seen in any of the *in-vivo* labelled data from this patient). *In-vivo* segmentations showed some other misalignment of both grey matter and SRLM labels, which can be seen in areas where neurons are visible in the histology without being obscured by grey matter labels. This likely reflects imperfect alignment between the *in-vivo* scan and *ex-vivo* histology, but may also be due to poor image quality in this patient making it difficult to correctly label hippocampal grey matter and SRLM.

**Figure 8.**
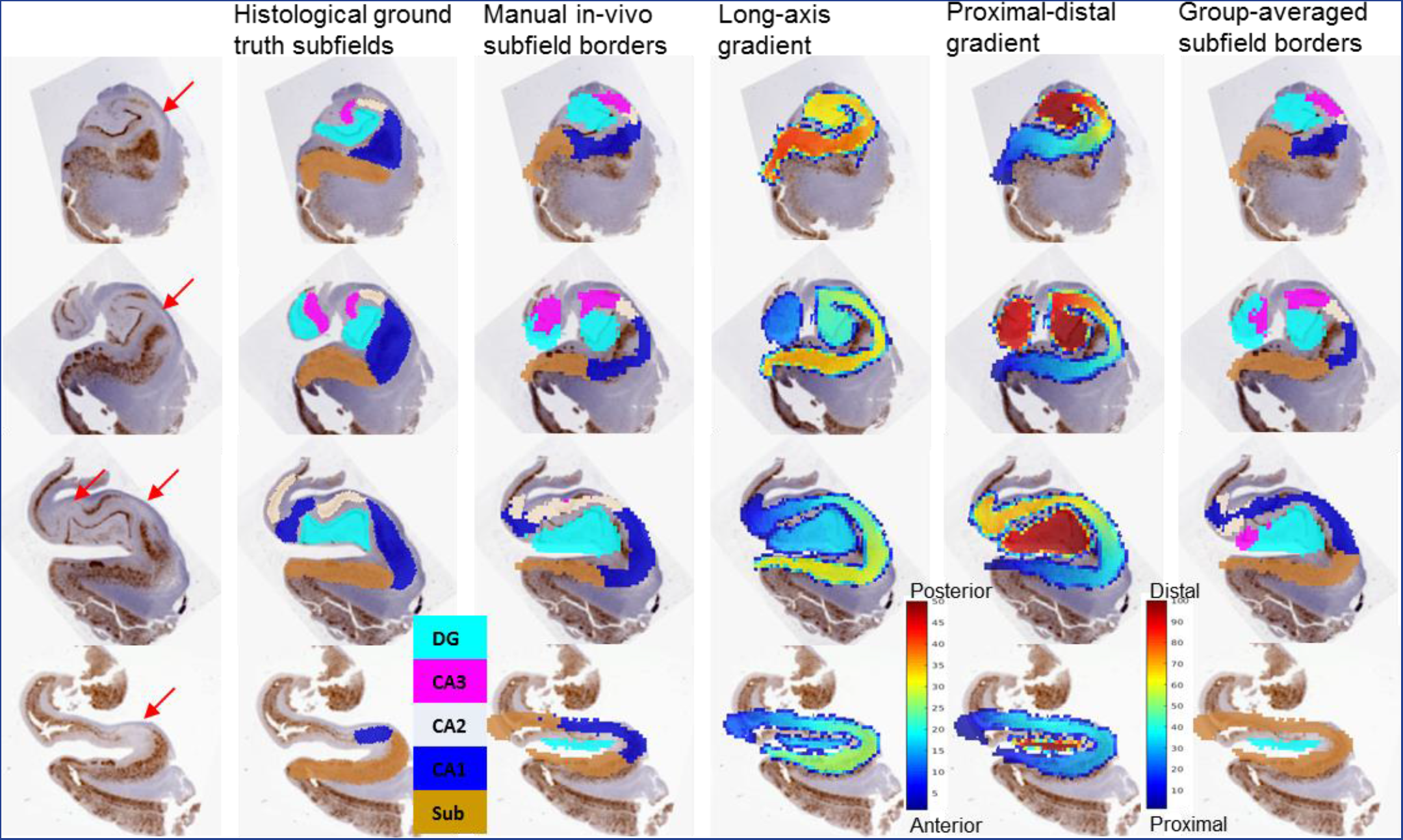
<double column width, coloured>. NeuN stain of resected hippocampal tissue with comparison of histologically segmented hippocampal subfields to *in-vivo* subfield labels and unfolding Laplace gradients in the same individual. Slices move from posterior (hippocampal body) to anterior through the hippocampal head, and are separated by 4.4mm. Red arrows indicated marked cell loss in distal CA1. The proximal-distal and long-axis gradients are surrounded by low color scaled voxels because of their interpolation when transforming to the histological space.

Some inconsistencies are noticeable in direct comparison between the *in-vivo* manual subfield segmentation (Figure 8 column 2) and the histological ground truth (Figure 8 column 3). This could be due to misalignment, tissue atrophy, poorer image quality, errors in the segmentation protocol, or inter-individual differences in subfield border locations. Note, however, that some key features are preserved in our segmentations: CA2 passes into plane twice in row 3 (i.e. appears in two different places), and both dentate gyrus and CA3 are seen in the uncus in row 2. Because these borders are curved medially in the head, they are difficult to capture in coronal slices alone and will vary drastically depending on the exact position of the slice.

The long-axis and proximal-distal gradient together make up our two-dimensional unfolded coordinate space, which can be examined in relationship to histology in Figure 8 columns 4 and 5. The long-axis gradient changes between coronal slices, as expected when moving from posterior to anterior, but also changes within each slice. This is because the anterior point is located in the most medial, vertical component of the uncus where each of the subfields has its natural terminus, rather than the absolute anterior of the hippocampus. Thus the long-axis gradient shows how any given coronal slice is out-of-plane with respect to the medial curvature of the uncus. The proximal-distal gradient identifies a set of potential subfield borders, which can also be adjusted depending on anterior-posterior extent. In row 3 this gradient can be seen to increase and then decrease as the gradient passes into and then out of the plane of view (i.e. from proximal to distal colours pass from green-yellow-orange-yellow-orange), similar to what is seen CA1 and CA2 in the corresponding histological ground truth images. We used this gradient in combination with the long-axis gradient to apply group-averaged subfield borders to this participant’s unfolded hippocampal space, which is shown in the far right column. This segmentation suffers from many of the same issues as the fully manual segmentation, but as in the manual segmentation, many of the key features of the hippocampal head are retained, including the passing into and out of plane for CA2 and the presence of CA3 in the uncus (Figure 8 rows 2 and 3, respectively). However, both in the unfolded group-average segmentation and in the fully manual segmentation, each of the *in-vivo* segmentation borders was incorrectly placed. Specifically, all subfield borders should be shifted more distally in rows 1 and 2, and in rows 3 and 4 the manually segmented border shifts are mixed whereas the group-averaged borders should be shifted more proximally. This resulted in no CA1 label the slice seen in row 4 of the unfolded group-average segmentation, despite its presence in the histological ground truth.

## 4 Discussion

Using isotropic T2 ultra-high resolution MR imaging, we were able to detect substructures of the hippocampus that can be leveraged to understand and quantify its complex and variable topology. Towards this end, we developed a methodology to ‘unfold’ or index this topology in a way that respects accounts of hippocampal subfields from the literature, inherently aligns tissues despite variable folding, and can be used to index or segment that tissue with a high level of precision and flexibility. Segmentations performed in the resulting unfolded space agree with quantitative metrics of intracortical myelin and capture key features shown in an *ex-vivo* validation sample from a patient with epilepsy.

### 4.1 Detection and labelling of the SRLM and hippocampal grey matter

High Dice Similarity Index was found for both hippocampal grey matter and SRLM using the segmentation instructions outlined in Supplementary Materials 1. This feature is critical in that it allows for the differentiation of folds throughout the entire hippocampus, and was necessary to constrain the proximal-distal Laplacian solution obtained here to the topology of hippocampal archicortex. Another constraint worth mentioning is that under our protocol, the SRLM is required to separate folds of hippocampal grey matter with no points of contact between the folds. This feature may be challenging to obtain in datasets generated with highly anisotropic acquisition, which will limit visibility of the SRLM in the medial extensions of the hippocampus, and also limit the rater’s ability to create a label with separated folds of hippocampal grey matter (as they may change too drastically between slices). However, with sufficiently thin slicing, it may still be possible to employ these criteria and procedures in anisotropic datasets.

### 4.2 Anatomical details of unfolding

The following list describes specific anatomical details that were correctly captured under our unfolded coordinate system:

- In the long-axis unfolding, a cross-section of hippocampal grey matter at an equipotential point in the hippocampal body reveals the same classic C-shaped orientation of grey matter as in coronal slices from histology and in extant MRI tracing protocols (Figure 4A, first right panel, red dotted line). Unlike in extant protocols based on coronal or other views, cross-sections in our unfolding at equipotential points in the hippocampal head also correctly reveal these C-shaped orientations of subfields (e.g. Figure 4A, second right panel, red dotted line).
- In the vertical component of the uncus, the C-shape of hippocampal grey matter flattens out to form a line (e.g. Figure 4A, third and fourth right panel, red dotted line). Here, the dentate gyrus passes most medially around the other subfields before extending upwards and reaching the vertical component of the uncus (Ding & Van Hoesen, 2015). This feature is accounted for in our unfolding by the manual placement of the ‘sink’ in the proximal-distal unfolding of hippocampal grey matter (Figure 4B, third and fourth right panels, turquoise bin; see also Supplementary Materials 1, step 3: labelling of extra-hippocampal structures).
- In the proximal-distal unfolding, the more proximal regions of hippocampal grey matter wrap around the absolute anterior of the hippocampus, moving from the inferior to superior side. This feature honors the descriptions of the subiculum provided by Ding & Van Hoesen (2015) and discussed in Introduction section 1.2 (Figure 4B, main model, green, blue, and yellow and cyan bins following dotted red arrow)

Limitations of current unfolded coordinate system:

- The ‘sink’ used for proximal-distal unfolding captures most of what corresponds to the granule cell layer of the dentate gyrus in histological studies. However, this tissue is so thin that it cannot easily be matched to the true granule cell layer seen in histology. Thus, we do not recommend using it as an independent region of interest for *in-vivo* MRI. Rather, we recommend combining it with area CA4, or the CA3 hilar region, as implemented in our manual subfield segmentations and in our unfolded subfield descriptions.
- The laminae of the dentate gyrus are known to be situated perpendicular to those of the other subfields (Duvernoy *et al.*, 2013). This is not respected by our laminar unfolding. Instead, most of the dentate gyrus is treated as being deep laminae (e.g. Figure 4C, second right panel, correct lamination shown in white dotted line). Thus, caution is necessary when the goal is to index the laminae of the CA4 and dentate gyrus.

Another strength of this ‘unfolded’ space pertains to the fact that all its distances are relative to the full size of the corresponding individual hippocampus, and can thus be applied across a range of hippocampal sizes and morphologies. With some adaptation of landmarks used as boundaries in the Laplace equation (i.e. source and sink), this protocol may also be applied to the characterization of abnormal hippocampi (due to abnormal development or neurological disease) or even those from other mammalian species.

### 4.3 Subfield borders in unfolded coordinate space

Figure 5 shows the mapping of subfields segmented in each participant’s native space to the standardized unfolded space. The fact that the SEM of the average unfolded borders was relatively low (i.e. accounts for a relatively small proportion of the area of each subfield) is surprising, given the large inter-individual variability of subfield locations in native space (e.g. highly variable digitations; see Ding & Van Hoesen, 2015). This result suggests that much of the variability in native space is due to differential curvature and folding of hippocampal tissue in development, rather than differences in the cytoarchitectural differentiation within this tissue.

Comparison of manual segmentations in native space to segmentations applied using the group-averaged borders in unfolded space revealed moderate to good spatial overlap, as determined by DSI, particularly when the smallest subfields CA2 and CA3 were combined (Figure 6). These DSI scores were also similar to manual segmentations. However, the sources of the remaining variability are not clear. They might reflect individual variability in subfield border locations that are not captured by our unfolded average borders. Alternatively, they might reflect deviations from the true subfield border locations in a manual segmentation in native space due to tracing errors.

### 4.4 Quantitative unfolded tissue properties

An important finding for the intracortical myelin estimates we obtained is that they appear to closely correspond to the average subfield borders used in our demonstration of subfield segmentations (Figure 7A). The subiculum and areas CA2 and CA3 appear to have greater myelin content than CA1 and the dentate gyrus, with the proximal part of the subicular complex showing greatest values. Though speculative, we suggest that this characteristic could reflect contributions of the perforant path passing through the subiculum, elevating myelin estimates due to the presence of white matter tracts. In area CA3, dense recurrent collaterals might contribute to elevated myelin estimates. An alternative explanation is that increased vasculature, which would also appear dark in T2-weighted images, contributes to this contrast. Support for this explanation comes from the observation that area CA2 is the most highly vascularized subfield in humans (Duvernoy *et al.*, 2013). Our findings also agree with those of Abraham *et al.* (2012) who examined intracortical myelin in histological samples, and those of Marques and Guetter (2013), who found similar differences in R1 MR intensities between the subfields.

Cortical thickness was also calculated using Laplacian streamlines and mapped in unfolded coordinate space (Figure 7B). Critically, these differences do not appear to correspond to the subfield borders. Note that thickness in the dentate gyrus was actually calculated perpendicular to the true laminae of the dentate gyrus because of the different orientation of tissue (see Section 3.2). Therefore, thickness in this region should not be considered to reflect true laminar structure. With this exception, our results are similar to thickness measures reported by Yushkevich *et al.* (2011), who found that thickness was highest between subiculum and CA1 and lowest in CA3, both in healthy individuals and those with mild cognitive impairment. However, these results differ from those reported by Burggren *et al.* (2008), who found that thickness was highest in CA3/dentate gyrus, particularly in the anterior hippocampus. This discrepancy may be due to the fact that their hippocampal unfolding did not account for the digitations in the hippocampal head, which could have led to overestimation of thicknesses.

Comparing both manual and unfolded group-average subfield segmentations performed in this study to intracortical myelin, we noted some discrepancies between border locations. In the example hippocampi examined, the subfield borders in the manual segmentation did not follow as smooth of a trajectory as in the unfolded segmentation (Figure 7C). This discrepancy could be due to individual differences in this participant’s subfield border locations that maynot be captured by the group average. However, we believe it is more likely due to limitations of the manual segmentation employed. For example, imperfect alignment between coronal slices in MRI with histological reference slices from Ding & Van Hoesen (2015) and Duvernoy *et al.* (2013) could cause subfield borders to shift, making them more jagged when in reality they follow a smooth trajectory. Detailed 3D histological examinations are needed to determine whether this is indeed the case. These issues may contribute to the variability observed between manual segmentations and unfolded group-averaged segmentations that are apparent in Figure 6. They also speak to more fundamental challenges related to the reliability and feasibility of manual segmentation protocols that are based primarily on coronal slices, which fall beyond the scope of the current paper. Nevertheless, the fact that these discrepancies largely average out in a sample of unfolded hippocampi, while still respecting differences in morphology, further highlight the strength of the approach presented here.

The unfolding of hippocampal grey matter accounts for much of the inter-individual variability related to differences in ontological folding. However, inter-individual differences in subfield border location beyond differences in folding structure (for example, that shown by Zeineh et al., 2015), as well as variability due to the presence of disease (as in the resected tissue examined here) may reflect additional sources of variability in unfolded subfield locations and size. This variability presents a significant challenge when aiming to apply unfolded group-average borders, as well as for manual or automated segmentation protocols that rely on geometric rules and structural landmarks. Thus, the use of other cues such as intracortical myelin or thickness may be useful in generating subject-specific subfield borders in future follow-up research.

### 4.5 Histological validation

Results from Figure 8 show some inconsistencies in our manual subfield segmentation as well as in our group-averaged unfolded space segmentations when compared to the histological ground truth in one resected tissue sample from a patient with epilepsy. In addition to general segmentation errors and natural inter-individual variability in border locations, some of these inconsistencies may arise because of tissue atrophy and poor grey matter labelling in this participant with a neurological condition as compared to healthy control participants. Such factors may make manual segmentation based on Ding & Van Hoesen (2015)’s descriptions, or the use of borders established in healthy control participants less appropriate for characterization of the hippocampi in a disease state. However, it should be noted that several features seen in the histology were still captured by our unfolding coordinate system, despite being absent from the subfield borders applied here. In particular, the proximal-distal gradient can be seen to increase and decrease along the length of a coronal slice, capturing how the CA1 passes into plane twice in the histological ground truth. Thus, although subfield borders may be shifted due to various factors, the coordinate space presented here still respects this feature of the hippocampal head. As such, this application illustrates how segmentations in unfolded coordinate space are able to capture critical structural complexities of the hippocampal head.

### 4.6 Hippocampal unfolding in the context of extant literature

One possible source of the recent controversy over hippocampal subfield borders relates to constraints that in order to be reliable, a coronal slice segmentation protocol has to make use of heuristics such as geometric rules with reference to visible intra-or extra-hippocampal landmarks. For these rules to be applicable across different hippocampal morphologies and MR image qualities, some level of simplification is necessary, reducing accuracy and precision. Given the large number of possible border locations, the unfolded coordinate system presented inherently allows for increased precision, even across hippocampi with varied morphologies, because it respects critical structural features of the hippocampal head without reliance on the heuristics mentioned. Thus, although we do not wish to present the specific subfield borders used here as an alternative to the efforts towards international harmonization by the Hippocampal Subfields Group, we hope that these efforts will include the structural considerations discussed in the current paper, and may also lead to exploration of methodologies other than manual segmentation. Furthermore, we anticipate that, once international consensus is reached, the resulting subfield borders can be applied using the unfolded coordinate system presented here, and be complemented by further characterization of inter-individual differences that can be captured with the present methodology. Given the increasing prevalence of high-resolution data in which the hippocampal SRLM can be identified throughout the length of the hippocampus, this appears to be a particularly promising avenue.

A final point worth noting is that the unfolded coordinate system offered here will also allow for easy implementation of further subfield divisions in future work. For example, Ding & Van Hoesen’s recent characterization of the hippocampal head (2015), as well as other histological evidence from humans and nonhuman animals (see Ding, 2013), reveal differentiation of the subiculum into distinct components, including the prosubiculum (postsubiculum in rodents), subiculum, presubiculum, and parasubiculum. Furthermore, some studies have documented functional differentiation between proximal and distal CA1 (Nakazawa *et al.*, 2016; Knierim *et al.*, 2014) and CA3 (Nakamura *et al.*, 2013). These findings highlight the increasing need for precision and standardization in indexing hippocampal tissue, as well as the need for flexibility in applying subfield labels so as to honour continuous new developments in tissue characterization. We believe that the unfolded coordinate system presented here can provide such a framework.

## 5 Conclusions

We have presented a new tool that promises to allow for *in vivo* characterization of the complex structure of the human hippocampal subfields in unprecedented detail. Manual segmentation with high anatomical detail poses many challenges for the generation of reliable protocols that are suitable for tracing of hippocampal subfields, in particular in the hippocampal head. However, considerations of regularities in hippocampal structure related to ontogeny offer ways in which computational tools, such as the Laplace equation, can be applied for indexing and segmenting hippocampal tissue in a way that preserves topology across individual difference. In the current study, we pursued an approach that took advantage of these considerations. Through computational unfolding of the hippocampus, the current protocol provides a coordinate system that can index hippocampal tissue in a precise and flexible manner, while capturing the noticeable inter-individual differences in morphology that have been documented in histological studies of this structure. This method critically depends on the visualization of the SRLM, or ‘crease’ along which the hippocampus is folded. We argue that this method offers several practical advantages over manual segmentation techniques. These advantages can be summarized as follows:

i. Unfolding hippocampal grey matter allows for indexing of analogous tissues (or sets of candidate boundary locations) across participants with variable morphologies.
ii. The unfolded coordinate space described can be used for inter-subject alignment and subsequent mapping of properties across the full long-axis and proximal-distal extents of the hippocampus, as illustrated here for intracortical myelin and cortical thickness measures.
iii. Segmentations applied in this unfolded coordinate space show good spatial overlap with, and may even correct for tracing errors in, the detailed manual subfield segmentations that were performed here. This coordinate system also captures subtle but critical structural features, as demonstrated in a direct comparison with a resected histological sample from a patient with epilepsy.

Future directions for this work include the integration of automatic tissue segmentation tools for detection of the SRLM, grey matter, and consideration of surrounding structures in order to reduce user input and improve reliability. Promising applications of this unfolded coordinate system include cross-species comparison and normative mapping of hippocampal tissue properties in health and disease.

## Acknowledgements

This project was supported by a CIHR Project Grant (Funding reference number: 148839) to AK and SK and by EpLink - The Epilepsy Research Program of the Ontario Brain Institute. Scanning was performed at Western’s Centre for Functional and Metabolic Mapping, supported by the Brain Canada, and the Canada First Research Excellence Fund to BrainsCAN.

Thanks to Dr. Robert Hammond for consultation on subfield border locations in histological samples.

